# TINC - a method to dissect transcriptional complexes at single locus resolution - reveals novel *Nanog* regulators in mouse embryonic stem cells

**DOI:** 10.1101/2020.04.03.023200

**Authors:** AS Knaupp, M Mohenska, MR Larcombe, E Ford, SM Lim, K Wong, J Chen, J Firas, C Huang, X Liu, T Nguyen, YBY Sun, ML Holmes, P Tripathi, FJ Rossello, J Schröder, CM Nefzger, PP Das, JJ Haigh, R Lister, RB Schittenhelm, JM Polo

## Abstract

Cellular identity is ultimately controlled by transcription factors (TFs), which bind to specific regulatory elements (REs) within the genome to regulate gene expression and cell fate changes. While recent advances in genome-wide epigenetic profiling techniques have significantly increased our understanding of which REs are utilized in which cell type, it remains largely unknown which TFs and cofactors interact with these REs to modulate gene expression. A major hurdle in dissecting the whole composition of a multi-protein complex formed at a specific RE is the shortage of appropriate techniques. We have developed a novel method termed TALE-mediated Isolation of Nuclear Chromatin (TINC). TINC utilizes epitope-tagged TALEs to isolate a specific genomic region from the mammalian genome and includes a nuclei isolation and chromatin enrichment step for increased specificity. Upon cross-linking of the cells and isolation of the chromatin, the target region is purified based on affinity purification of the TALE and associated nucleic acid and protein molecules can be subjected to further analyses. A key TF in the pluripotency network and therefore in embryonic stem cells (ESCs) is NANOG. It is currently not fully understood how *Nanog* expression is regulated and consequently it remains unclear how the ESC state is maintained. Using TINC we dissected the protein complex formed at the *Nanog* promoter in mouse ESCs and identified many known and numerous novel factors.

## Introduction

Pluripotent stem cells (PSCs) carry immense therapeutic potential due to their ability to give rise to any cell type of the body and their capacity to self-renew indefinitely. The most common *in vitro* models of PSCs are embryonic stem cells (ESCs), which are obtained from the inner cell mass (ICM) of the blastocyst (Martin 1981; Kaufman et al. 1983) and induced pluripotent stem cells (iPSCs), which can be derived from somatic cells through overexpression of the four transcription factors (TFs) OCT4, SOX2, KLF4 and C-MYC (OSKM) (Takahashi and Yamanaka 2006). Besides OSKM, the homeobox TF NANOG has been shown to be involved in pluripotency maintenance of the ICM and of ESCs/iPSCs (Mitsui et al. 2003; Jose Silva et al. 2009). Together with OCT4 and SOX2, NANOG forms the core transcriptional regulatory circuitry that drives the ESC state (Boyer et al. 2005). These three TFs co-occupy many targets and collaborate to activate the expression of self-renewal and pluripotency genes including themselves and to suppress the expression of genes that promote differentiation (Loh et al. 2006; Boyer et al. 2005; Chen et al. 2008; Marson et al. 2008; Kim et al. 2008). Interestingly, out of the three core pluripotency TFs only NANOG seems to have the capacity to mediate LIF/STAT3-independent self-renewal of ESCs (Mitsui et al. 2003; Chambers et al. 2003).

Besides pluripotency maintenance, NANOG plays an important role during cellular reprogramming (José Silva et al. 2006; Jose Silva et al. 2009; Yu et al. 2007). There is evidence, however, that somatic cells can be reprogrammed in the absence of NANOG, albeit at much lower efficiency (Carter et al. 2014). Additionally, there is evidence that ESCs can be maintained upon genetic deletion of *Nanog* although the cells are prone to differentiation (Chambers et al. 2007). This observation, together with work revealing that *Nanog* is expressed in a heterogeneous manner in ESCs (Chambers et al. 2007; Kalmar et al. 2009; Kumar et al. 2014; Abranches et al. 2014), suggests that the pluripotency network can cope with variable levels of NANOG. Fluctuations in NANOG levels seem to allow ESCs to regulate lineage commitment, with high NANOG levels impeding and low levels promoting response to differentiation cues (Kalmar et al. 2009; Abranches et al. 2014; Chambers et al. 2003). How the expression of *Nanog* is regulated in ESCs remains, however, still not fully understood.

As any given gene, the expression of *Nanog* is controlled by TFs, which bind to specific regulatory elements (REs) and recruit other factors such as TFs, cofactors and epigenetic regulators to modulate gene expression. There are at least two major REs controlling *Nanog* expression in ESCs: one in the promoter region and an enhancer approximately 5 kb upstream of the TSS that loops to interact with the *Nanog* promoter (Kagey et al. 2010; Apostolou et al. 2013). NANOG has been shown to bind to both of these REs (Chen et al. 2008; Kim et al. 2008), and this NANOG activity has been associated with auto-activation (Q. Wu et al. 2006) as well as auto-repression (Fidalgo et al. 2012; Navarro et al. 2012). These positive and negative *Nanog* autoregulatory feedback loops are linked to NANOG interacting with different TFs. For example, SALL4 has been shown to physically associate with NANOG to activate *Nanog* expression (Q. Wu et al. 2006), whereas *Nanog* auto-repression has been shown to depend on its interaction with ZFP281, which in turn recruits the NuRD repressor complex (Fidalgo et al. 2012). Furthermore, positive transcriptional regulation of *Nanog* has been associated with binding of OCT4 and SOX2 to the *Nanog* promoter (Rodda et al. 2005), whereas *Nanog* auto-repression has been shown to be independent of OCT4 and SOX2 (Navarro et al. 2012). This suggests that fine-tuning of *Nanog* expression levels might rely on differences in TF and cofactor interactions. However, while some of the proteins that interact with these RE are known, the full characterization of the complexes formed at any of these REs remains largely elusive.

One major reason for this lack of knowledge is that dissecting the whole molecular composition of a regulatory complex formed at a specific RE is very difficult to achieve. Chromatin immunoprecipitations (ChIPs) have been invaluable to study the genome-wide DNA binding events of proteins of interest, however, they require *a priori* knowledge in order to select a candidate protein, antibodies appropriate for this application and they allow the interrogation of only one protein at the time (reviewed in (Furey 2012; Collas 2010)). Consequently, being able to simultaneously analyze the entire set of proteins that interact at a specific RE would be advantageous, however, this is extremely challenging for two main reasons: 1) a very low abundant genomic region (e.g. two copies per cell) has to be specifically targeted and 2) the interacting proteins have to be enriched efficiently to be detected. Unlike DNA and RNA molecules, which can be easily amplified (e.g. PCR), proteins have to be interrogated without further amplification, which poses a significant challenge for low abundant RE-associated proteins. High-resolution liquid chromatography tandem mass spectrometry (LC-MS/MS) is the method of choice for such analyses. Hence, the number of proteins, which can be identified and quantified at a specific genomic locus, is primarily governed by the intrinsic properties of the mass spectrometer such as mass accuracy, resolving power and sensitivity, which in turn dictate the lower limit of detection and quantification for individual species (typically in the femtomole-on-column range).

Several groups have set out to overcome these limitations by developing various locus-specific isolation or proximity labelling methods in combination with mass spectrometry. These include nucleic acid hybridization-based approaches (Ide and Dejardin 2015; Déjardin and Kingston 2009; Antão et al. 2012; Kennedy-Darling et al. 2014) and approaches that utilize DNA-binding proteins including LexA (Byrum et al. 2012; Fujita and Fujii 2011), TetR (Pourfarzad et al. 2013), Cas9 (Waldrip et al. 2014; Fujita and Fujii 2013; X. Liu et al. 2017; Gao et al. 2018; Schmidtmann et al. 2016; Tsui et al. 2018; Qiu et al. 2019) or TALE proteins (Fujita et al. 2013; Byrum, Taverna, and Tackett 2013; Fang et al. 2018). Although nucleic acid hybridization-based approaches pioneered the field, they require intensive probe-specific optimizations (Ide and Dejardin 2015; Déjardin and Kingston 2009; Antão et al. 2012), which may account for why this approach has not yet been widely adopted nor translated to single copy elements in mammalian cells. To target such challenging genomic regions in the mammalian genome, TALE (Fang et al. 2018) and CRISPR/Cas9 (Waldrip et al. 2014; Fujita and Fujii 2013; X. Liu et al. 2017; Gao et al. 2018; Schmidtmann et al. 2016; Tsui et al. 2018; Qiu et al. 2019) approaches are starting to emerge as the systems of choice as, unlike TetR and LexA proteins, they do not rely on genetic engineering of binding sites into the target sequence.

Here, we describe a novel epigenetic method that allows the isolation of a specific genomic region from the complex mammalian genome termed TALE-mediated Isolation of Nuclear Chromatin (TINC). TINC utilizes an epitope-tagged TALE in combination with a nuclei isolation and chromatin enrichment step based on Kustatscher et al., 2014 to further increase the signal-to-noise ratio. Even though the only other mammalian cell-validated TALE-based single locus isolation technique utilizes whole cell lysate (Fang et al. 2018) we chose to include these two additional purification steps (nuclei and chromatin) to reduce cytoplasmic and free nuclear protein contaminant contribution. Using TINC, we interrogated the protein complex formed at the *Nanog* promoter in mouse ESCs. We were able to not only detect many proteins previously shown to interact with this key pluripotency RE such as OCT4, SOX2 and NANOG (Chen et al. 2008; Kim et al. 2008) but more importantly also many novel candidates.

## Results

### Development of TINC to target the *Nanog* promoter in mESCs

In order to interrogate the protein complex formed at the *Nanog* promoter in ESCs, we developed TINC (Figure 1A); an epigenetic technique applicable to essentially any genomic region. In short, a construct driving the expression of an epitope tagged TALE (3xHA-tag) designed to bind to the sequence of interest is introduced into mammalian cells (e.g. via transfection/transduction). Upon confirmation that the TALE is expressed (e.g. via western blot/immunofluorescence), the cells are fixed with formaldehyde, nuclei isolated followed by chromatin isolation (Kustatscher et al. 2014) to enrich chromatin-associated proteins and to further decrease background noise. The chromatin is then sheared by sonication and the target region is isolated based on affinity purification of the TALE (e.g. utilizing anti-HA agarose). After elution and further purification, nucleic acids and proteins can then be analysed by appropriate techniques including quantitative polymerase chain reaction (qPCR) and LC-MS/MS, respectively. This allows the assessment whether the genomic locus of interest has been isolated and which proteins are associated with this genomic locus.

**Figure 1.**
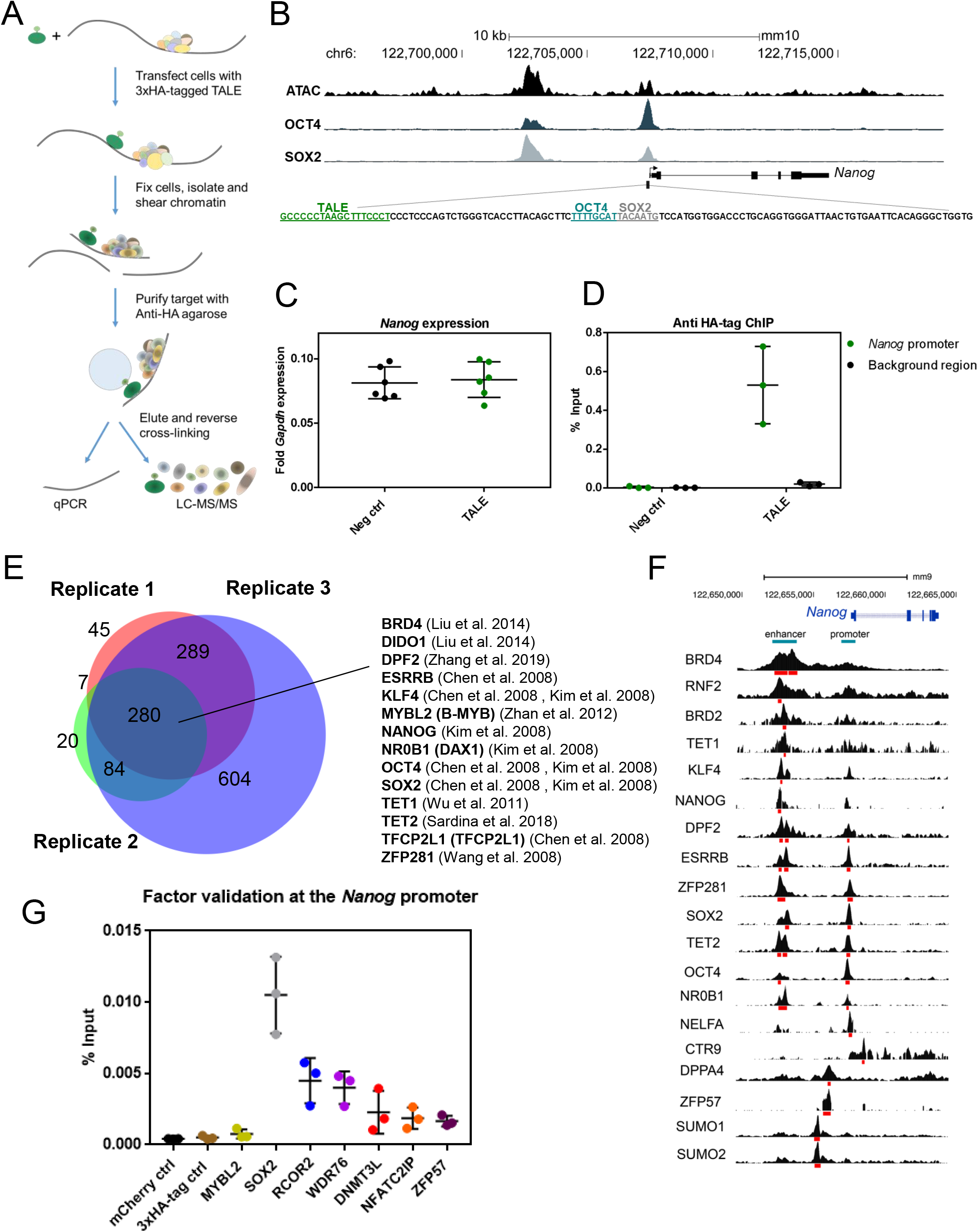
TINC allows isolation of the Nanog promoter complex from mouse ESCs. (A) Schematic of the TINC method. (B) A TALE was designed to bind to the proximal promoter of *Nanog* in mouse ESCs upstream of the OCT4 and SOX2 binding sites (Rodda et al. 2005). This region is located in open chromatin as shown by ATAC sequencing, which also shows the *Nanog* enhancer located approximately 5kb upstream of the TSS and which is also bound by OCT4 and SOX2 (Knaupp et al. 2017). The TALE binding site is indicated in green, the OCT4 binding site in blue and the SOX2 binding site in gray. (C) qPCR revealed that the expression of the TALE does not alter the expression of *Nanog*. *Nanog* expression levels were normalized to the housekeeping gene *Gapdh*. (D) HA-tag ChIP-qPCR confirmed enrichment of the *Nanog* promoter only in TALE expressing cells and not in empty vector transfected control cells. This enrichment is specific to the *Nanog* promoter as a background genomic region (e.g. *Sox2* regulatory region 2) was not enriched by the TALE. (E) Overlap of the proteins identified in TINC replicates 1, 2 and 3. The 280 proteins identified by all three TINC replicates contain many of the previously published binders of the *Nanog* promoter and were hence used for further analysis. (F) Validation of proteins identified by TINC utilizing publically available ChIP-seq data sets in the database ChIP-Atlas (Oki et al. 2018). Below each sample’s BigWig track (black) are the respective peaks called by ChIP-Atlas (red). (G) ChIP-qPCR of novel *Nanog* promoter binders identified by TINC. MYBL2 and SOX2 were used as positive controls.

To investigate the protein complex regulating the transcription of the master pluripotency regulator NANOG, TINC was utilized to isolate the *Nanog* promoter from mouse ESCs. A TALE was designed to bind upstream of the OCT4 and SOX2 binding sites (Rodda et al. 2005) in this key RE (Figure 1B) and a clonal ESC line stably expressing this TALE was created (Figure S1A). A negative control line, transfected with the empty vector, was also generated (Figure S1A). TALE-expressing ESCs were characterized by unaltered *Nanog* expression (Figure 1C) and importantly, ChIP of the TALE resulted in clear enrichment of its genomic target: the *Nanog* promoter (Figure 1D). Furthermore, the capacity to give rise to cells from all three germ layers in teratoma assays was unaffected (Figure S1B). Altogether, these results suggest that binding of the TALE to the *Nanog* promoter does not change expression of the target gene nor the pluripotency potential of the cells.

To determine the proteins that interact with the *Nanog* promoter in mouse ESCs, we performed TINC and the enriched proteins were analysed by LC-MS/MS (Figures 1E and S1C). The proteins that were also detected in the negative controls were considered as background (please see discussion) and consequently discarded from further analysis (Figure S1C). Some of these proteins were found to be enriched in the TALE samples in comparison to the negative controls (data not shown) suggesting they might also interact with the *Nanog* promoter, however, we decided to be stringent and rather increase the number of false negatives than of false positives. To further increase the accuracy of our results, we included an additional cut-off (Vermeulen and Déjardin 2020) and only utilized TINC replicates for further analysis that identified the three expected key pluripotency regulators OCT4, SOX2 and NANOG. Overlap of the proteins detected uniquely in the TALE samples in three TINC replicates revealed 280 candidates to be present in all three replicates (Figure 1E). Importantly, besides OCT4, SOX2 and NANOG these included many other proteins that had previously been shown to interact with the *Nanog* promoter in mouse ESCs such as BRD4 (W. Liu et al. 2014), DIDO1 (Y. Liu et al. 2014), DPF2 (Zhang et al. 2019), ESRRB (Chen et al. 2008), KLF4 (Chen et al. 2008; Kim et al. 2008), MYBL2 (also known as B-MYB) (Zhan et al. 2012), NR0B1 (also known as DAX1) (Kim et al. 2008), TET1 (H. Wu et al. 2011), TET2 (Sardina et al. 2018), TCFCP2L1 (also known as TFCP2L1) (Chen et al. 2008) and ZFP281 (Wang et al. 2008). To further confirm the validity of our TINC technique, we utilized the database ChIP-Atlas, which allows data mining and visualization of published chromatin immunoprecipitations (ChIP) sequencing data sets (Oki et al. 2018). ChIP-Atlas revealed binding of an additional eight proteins within this genomic region in ESCs: BRD2, CTR9, DPPA4, NELFA, RNF2, SUMO1, SUMO2 and ZFP57 (Figure 1F). Interestingly, these proteins seem to interact with three distinct sites: the *Nanog* promoter and/ or the −5kb enhancer or a region in between these two REs.

We also validated by ChIP-qPCR five novel candidates, which are upregulated during MEF reprogramming (Knaupp et al. 2017) but had previously not been shown to interact with the *Nanog* promoter: RCOR2 (REST corepressor 2), DNMT3L (DNA (cytosine-5)-methyltransferase 3-like), NFATC2IP (NFATC2-interacting protein), WDR76 (WD repeat-containing protein 76) and ZFP57 (zinc finger protein 57) (Figure 1G). We included ZFP57 for which the ChIP-Atlas data base suggested targeting of the *Nanog* promoter (Figure 1F) while the original publication did not call this binding site (Strogantsev et al. 2015). To overcome the lack of ChIP antibodies against most of these proteins, we expressed 3xHA-tagged versions of these candidates in ESCs as well as of SOX2 (Chen et al. 2008; Kim et al. 2008) and MYBL2 (Zhan et al. 2012) as positive controls. ChIP-qPCR confirmed enrichment of these proteins at the *Nanog* promoter, with enrichment values in between the two positive controls SOX2 and MYBL2, the latter of which showed only very poor enrichment (Figure 1G).

Altogether, this demonstrates that TINC is an appropriate technique for the interrogation of the protein complex formed at the *Nanog* promoter in ESCs, and hence any given RE in the mammalian genome, with the potential to discover hundreds of interacting proteins.

### Interrogation of the protein complex formed at the *Nanog* promoter using TINC

The 280 proteins found to interact at the *Nanog* promoter were further characterized to unveil their potential roles at this RE. Unsurprisingly, protein class ontology analysis showed the strongest enrichment for nucleic acid binding proteins (Figure 2A). Additionally, protein modifying enzymes (e.g. non-receptor serine/threonine protein kinases, proteases, protein phosphatases and ubiquitin-protein ligases), gene-specific transcriptional regulators and chromatin/chromatin-binding or chromatin-regulatory proteins were found to be strongly enriched (Figure 2A). Importantly, analysis of the 57 transcription factors identified by TINC revealed that they were enriched for biological processes and KEGG pathways associated with stem cell maintenance and pluripotency (Figure 2B). Further characterization of all 91 epigenetic modifiers showed that besides TFs associated with transcriptional activation and repression, many were histone modification writers or their cofactors (Figure S2A). Only a minority of factors we identified were chromatin remodelling (co-) factors or DNA modifiers (e.g. mediating DNA hydroxymethylation). Consequently, it is unsurprising that while the molecular target of the TFs is DNA, the molecular target of the majority of the other epigenetic modifiers we identified at the *Nanog* promoter are histones (Figure S2A). These results suggest that various and different epigenetic modifiers are required to regulate *Nanog* expression and that they exert their roles at various levels including modifications of the DNA and histone landscape.

**Figure 2.**
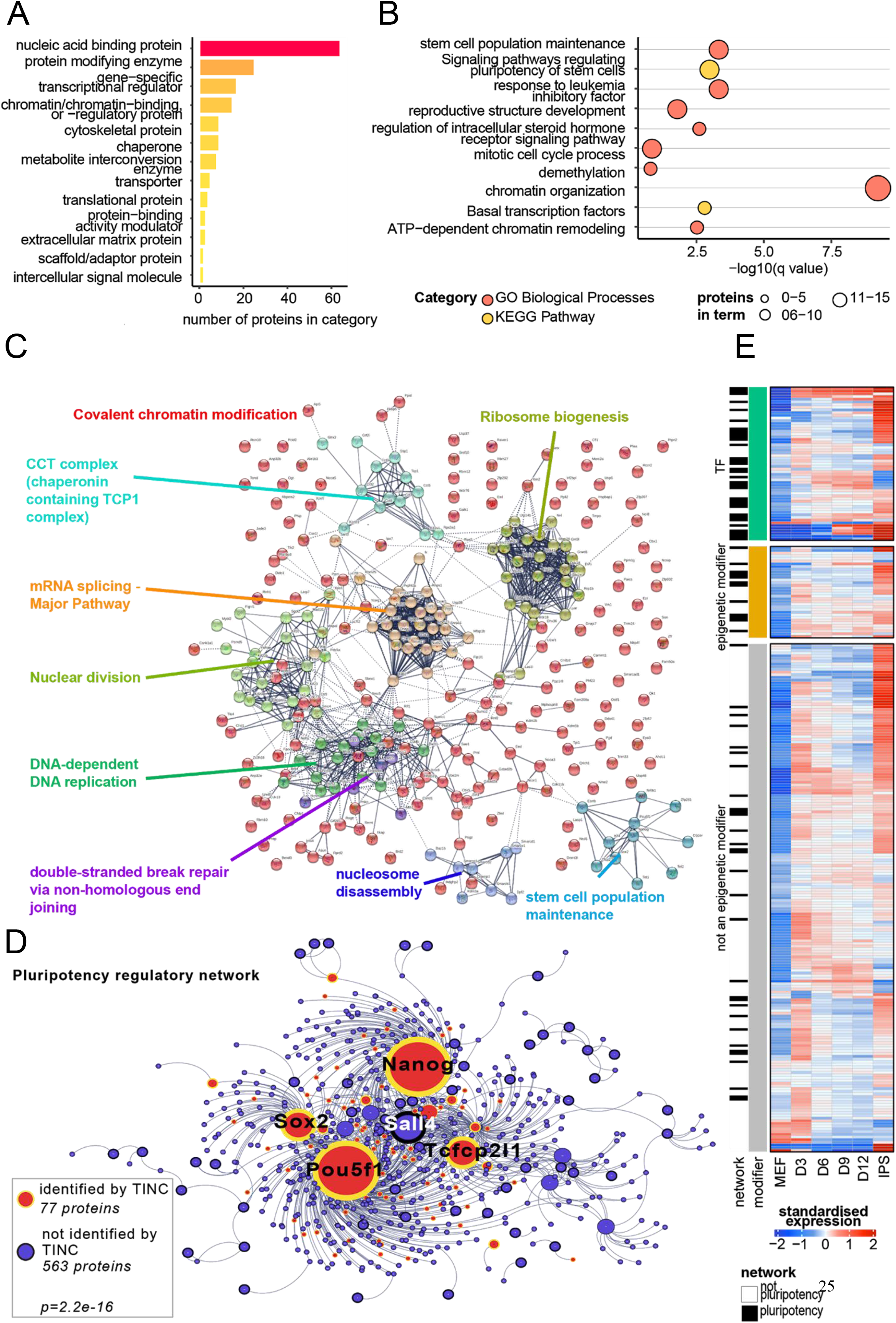
Characterization of proteins interacting with the *Nanog* promoter. (A) Protein class categories for all 280 proteins identified by TINC. The color gradient corresponds to the number of proteins in the term with yellow being a few, and red being many. (B) GO Biological Process and KEGG pathway enrichment of transcription factors identified by TINC. (C) Protein-protein interaction enrichment of the TINC proteins and k-means clustering analysis of the identified network by STRING as well as biological processes enriched for each of the clusters. (D) The pluripotency regulatory network overlapped with proteins identified by TINC at the *Nanog* promoter. Significant enrichment of the TINC proteins in the pluripotency regulatory network was identified by a Fisher’s exact test. (E) Proteins identified by TINC with standardized expression values throughout the reprogramming process from MEFs to iPSCs. Proteins have been categorized into TFs, other epigenetic modifiers and all additional proteins. The black bars indicate whether they are a part of the pluripotency network from Figure 2D.

In order to gain further insight into the protein interaction network we conducted protein network analysis and k-means clustering of all proteins identified by TINC using STRING (Szklarczyk et al. 2019). This approach revealed nine distinct clusters associated with covalent chromatin modifications, mRNA splicing, ribosome biogenesis, nuclear division, DNA replication, CCT (chaperonin containing TCP1 complex), stem cell population maintenance, nucleosome disassembly and double-strand break repair (Figure 2C). This further supports the above-mentioned notion that the proteins regulating the transcription of *Nanog* are required to exhibit differing functions at the DNA, RNA and protein level.

Since NANOG is one of the master regulators of pluripotency, we next sought to determine whether other key regulators of the pluripotency network control the transcription of this gene. Comparison of the proteins we identified at the *Nanog* promoter with the previously defined pluripotency regulatory network (Nefzger et al. 2017; Xu et al. 2013, 2014) revealed a significant enrichment of 77 proteins including OCT4, SOX2, NANOG and TCFCP2L1 (Note: SALL4 was identified in two of the three TINC replicates) (Figure 2D). This indicates that over a tenth of the key regulators of the pluripotency network are required to control the transcription of *Nanog*. These 77 proteins are broadly classified as nucleic acid binding proteins, transcriptional regulators, chromatin-chromatin binding or regulatory proteins, chaperones and protein modifying enzymes (phosphatases, proteases, kinases and E3 ubiquitin ligases) (Figure S2B).

As mentioned previously, NANOG does not only play an important role during pluripotency maintenance but also in attaining the pluripotent state during reprogramming (José Silva et al. 2006; Jose Silva et al. 2009; Yu et al. 2007). We therefore determined the expression dynamics during reprogramming of the proteins associated with the *Nanog* transcriptional complex and hence, when the pluripotency-associated complex is potentially established during this cellular conversion (Figure 2E). Analysis of the expression of the 280 TINC proteins during reprogramming of mouse embryonic fibroblasts (MEFs) into iPSCs revealed that 192 of these proteins are upregulated during this cellular conversion (Figure 2E). This suggests that these 192 proteins are specific to pluripotency meanwhile the remaining 88 proteins play common roles across cell types. *Nanog* expression is maintained at low levels during reprogramming and only increases significantly between day 12 and the iPSC state (Figure S2C). In order to determine whether this change in *Nanog* expression might be due to a change in the expression of some of the proteins that regulate the expression of *Nanog* in the pluripotent state we correlated their expression to the expression of *Nanog* throughout the reprogramming process (Figures S2D and S2E). Interestingly, most of these proteins were upregulated at the later stages of reprogramming and the expression of most of these proteins was positively correlated with the expression of *Nanog* (Figure S2D). 16% of these highly correlated genes are TFs, 10% are epigenetic modifiers and 51% are other proteins many of which are associated with RNA modification and splicing. Furthermore, the 25% most correlated genes are associated with chromatin organization, dedifferentiation, cell cycle and genetic imprinting development (Figure S2F). Intriguingly, all of the proteins that were negatively correlated with the expression of *Nanog* were transiently upregulated during the early stages of reprogramming (Figure S2E). These genes are mostly involved in mRNA processing and protein ubiquitination (Figure S2F). Overall, this indicates that many of the proteins identified at the *Nanog* promoter in the pluripotent state are upregulated only at the final stages of cellular reprogramming. Importantly, this coincides with the initiation of *Nanog* expression during this cellular conversion (Figure S2C).

## Discussion

Here, we describe TALE-mediated Isolation of Nuclear Chromatin (TINC), a novel epigenetic technique that allows isolation of a single genomic locus from mammalian cells for further molecular interrogation. TINC utilizes epitope tagged TALE proteins, which have previously been shown to possess the capacity to enrich a single RE from mammalian cells (Fang et al. 2018). In order to minimize background contribution from cytoplasmic contaminants as well as from proteins specifically enriched by the TALE but without target locus-related functions (e.g. proteins that interact with the nascent TALE polypeptide chain during translation), TINC contains a nuclei isolation and a chromatin enrichment step (Kustatscher et al. 2014).

We used TINC to interrogate the regulatory complex formed at the *Nanog* promoter in ESCs and only considered proteins that were solely detected in the TALE samples but not the negative controls. We acknowledge that this approach might discard some proteins that are indeed part of the studied complex (e.g. proteins that are enriched in the TALE samples in comparison to the negative controls), that therefore become false negatives. Additionally, Jérôme Déjardin, who, together with Robert Kingston, published the first protocol for the isolation of specific genomic DNA segments (human telomeres), stated recently that extreme caution should be exerted when interpreting such locus specific data that did not recover expected proteins (Vermeulen and Déjardin 2020). In agreement with Vermeulen and Déjardin, 2020, we therefore utilized previously identified binders as positive controls (e.g. the master pluripotency regulators OCT4, SOX2 and NANOG) and excluded one TINC replicate that did not allow identification of SOX2. Furthermore, we only considered proteins identified in all three valid TINC replicates as bound to the *Nanog* promoter. This stringent approach allowed us to identify many previously published *Nanog* promoter interactors including the key pluripotency factors OCT4, SOX2 and NANOG (Chen et al. 2008; Kim et al. 2008) as well as many proteins that had previously not been shown to bind to this RE. However, even if we had taken a completely unbiased approach and had not excluded the fourth TINC replicate that was missing SOX2, we would have identified many known *Nanog* binders besides OCT4 and NANOG including BRD4, DIDO1, DPF2, KLF4, MYBL2, NR0B1, TET1, TET2 and TFCP2L1 and more than 100 novel candidates. This suggests that while known binders are very helpful for setting analysis cut-offs, TINC can also give significant insight into REs without prior knowledge on interacting proteins.

To support the validity of our TINC method we confirmed by conventional ChIP-qPCR the binding of five novel proteins: RCOR2, ZFP57, DNMT3L, NFATC2IP and WDR76 to the *Nanog* promoter. As mentioned above, we needed to tag these proteins since we could not source ChIP antibodies. It is possible that the epitope tag interfered with folding or binding of these TFs and that is why ChIP could not enrich MYBL2 well at this RE. However, the commercial antibody previously used to show MYBL2 binding to the *Nanog* promoter (Zhan et al. 2012) has been discontinued. On that note, the authors of this study tested five commercial antibodies and only one of them showed binding of MYBL2 at known target genes (Zhan et al. 2012). Altogether, this highlights the shortcoming of conventional ChIP and why an unbiased method such as TINC, which does not rely on specific antibodies, is of major importance for many scientific fields. Nevertheless, our validation experiments confirmed *Nanog* promoter binding of these novel factors, some of which, including RCOR2, had previously been identified to play a role in pluripotency. RCOR2 is a subunit of the histone demethylase LSD1 complex in mouse ESCs and it has been shown to be able to replace SOX2 during somatic cell reprogramming into iPSCs (Yang et al. 2011). The authors of this work also showed that knockdown of *Rcor2* in ESCs results in significantly decreased expression levels of *Nanog* and *Sox2*. Based on RCOR2 ChIP-qPCR for selected differentiation marker genes, the authors proposed an indirect mechanism for the decrease of these pluripotency genes upon RCOR2 depletion. However, our TINC and ChIP data suggest that *Nanog* is a direct target of RCOR2. The observed decrease in *Nanog* expression upon RCOR2 depletion (Yang et al. 2011) suggests that RCOR2 acts as transcriptional activator of this pluripotency gene. Similarly, knockout of *Zfp57*, a transcriptional regulator conventionally associated with maintaining maternal and paternal genomic imprinting, has previously been associated with transcriptional deregulation of 146 genes in mouse ESCs including *Nanog (Riso et al. 2016)*. While the authors did not detect ZFP57 binding within 100 kb of *Nanog* and hence did not consider it as one of the genes directly regulated by this TF (Riso et al. 2016) our TINC and ChIP data indicate that *Nanog* is a direct target of ZFP57. On the other hand, it has previously been shown that knockdown of DNMT3L, a catalytically inactive member of the DNMT3 family of *de novo* DNA methyltransferases, does not alter the expression of *Nanog* in mouse ESCs but their differentiation potential (Neri et al. 2013). Together with our data, which revealed binding of this epigenetic modulator at the *Nanog* promoter in ESCs, this suggests that DNMT3L might play a role in the silencing of *Nanog* during differentiation; in particular into primordial germinal cells (Neri et al. 2013). Even though the expression of NFATC2IP and WDR7 increases significantly during reprogramming of MEFs into iPSCs (Knaupp et al. 2017), their roles in ESCs remain largely unknown. While NFATC2IP has been associated with transcriptional regulation of cytokine genes in T helper cells through methylation of histone H4 at arginine 3 (H4R3) in specific cytokine promoters (Fathman et al. 2010), WDR76 might play a role in epigenetic transcriptional regulation as it preferentially binds to 5-hydroxymethylcytosine (Spruijt et al. 2013). Our work now suggests that both of these proteins might also play a role in *Nanog* regulation as we showed (via TINC and ChIP-qPCR) that NFATC2IP and WDR76 associate with its promoter.

Overall, TINC allowed us to identify a large number of proteins associated with functions that are connected to *Nanog* transcriptional regulation such as stem cell population maintenance, covalent chromatin modifications, nucleosome disassembly and mRNA splicing. There is evidence that pre-mRNA splicing can occur post-transcriptionally but most splicing events seem to take place during the process of transcription (reviewed in (Brugiolo, Herzel, and Neugebauer 2013)). Our data supports this concept, since we detected several proteins associated with mRNA splicing at the *Nanog* promoter. Besides proteins related to transcriptional regulation, we found many proteins associated with nuclear division, DNA replication and double strand break repair some of which might be explained by the fact that the cells were potentially at many different stages of the cell cycle at the point of analysis, as ESCs are self-renewing and constantly proliferating (Lobanok et al. 2009). Additionally, our TINC data provides plausible explanations for several previously published observations. For example, Loh and colleagues showed that depletion of the histone demethylase JMJD1A (also known as KDM3A) in mouse ESCs leads to a significant decrease in *Nanog* expression (Loh et al. 2007). The authors did not detect JMJD1A at the *Nanog* promoter in ChIP experiments and consequently attributed this change in *Nanog* expression to an indirect effect. Our TINC analysis, however, revealed that the *Nanog* promoter is indeed a direct target of JMJD1A hence suggesting JMJD1A might play a role in histone H3 lysine 9 (H3K9) demethylation in this region.

Furthermore, Gingold and colleagues performed a genome-wide RNAi screen using a *Nanog*-GFP reporter mouse ESC line under differentiation-promoting conditions to identify factors required for pluripotency maintenance and differentiation (Gingold et al. 2014). Interestingly, we identified 234 of their candidates at the *Nanog* promoter but only 13 of them were associated with a change in *Nanog* expression one of which is *Zfp57 (Gingold et al. 2014)*. This suggests that the majority of the proteins that interact with the *Nanog* promoter are functionally redundant under the conditions tested. As to whether they may play a role during differentiation into cell types not promoted by the author’s mild retinoic acid induced-differentiation conditions, remains elusive. Also, it should be taken into consideration that a change in *Nanog* expression below the threshold set by the authors (e.g. |Z-score|>2) might still have a significant effect on the cells’ self-renewal and/or pluripotency potential. Out of the 13 proteins associated with a change in *Nanog* expression, knockdown of seven led to an increase (*Blm*, *Ik*, *Pogz*, *Pola2*, *Ppil2*, *Ppp1ca* and *Zfp57*) whereas six (*Brd4*, *Cbx1*, *Chek2*, *Fen1*, *Rnh1* and *Wiz*) led to a decrease in *Nanog*-GFP fluorescence (Gingold et al. 2014). This suggests that transcriptional activators and repressors co-occupy the *Nanog* promoter. It has previously been shown that some of the *Nanog* repressors that were found to interact with this RE in ESCs including ZFP281 (Fidalgo et al. 2011) and TCF3 (Yi, Pereira, and Merrill 2008) play a role during differentiation by helping the cell to exit the pluripotent state. Consequently, integration of our TINC data with this RNAi screen potentially suggests that the *Nanog* promoter is occupied by a large number of proteins whose primary role is not to maintain the pluripotent state but rather to allow exit of this state upon exposure of the cell to specific differentiation cues.

In summary, TINC allowed us to interrogate the regulatory complex formed at the *Nanog* promoter in an unbiased manner and provided insight into general molecular processes of the cell. In regards to *Nanog*, our data suggest transcriptional regulation of this key pluripotency TF occurs at many different levels since factors with various functions including DNA modifiers, histone modifiers, chromatin remodelers, transcriptional activators and transcriptional repressors reside at its promoter. Together, this implies a highly complex and coordinated interplay of these factors mediating fine-tuning of the expression of *Nanog*. Furthermore, our data suggest that many factors that aid in downregulation of *Nanog* during differentiation already reside at its promoter in the pluripotent state.

## Materials and Methods

### Cell culture

ESCs were cultured in gelatin coated dishes at 37°C, 5% CO_2_ in ESC medium (KnockOutTM DMEM (Life Technologies) containing 15% Fetal Bovine Serum (Thermo Scientific), GlutaMAX Supplement (Life Technologies), MEM Non-Essential Amino Acids Solution (Life Technologies), Penicillin-Streptomycin (Life Technologies), 0.1% (v/v) 2-Mercaptoethanol (Life Technologies) and 1000 U/mL Leukemia Inhibitory Factor (LIF) (Merck Millipore)) supplemented with 2i (1mM PD0325901 (StemMACS) and 3mM CHIR99021 (StemMACS)). To maintain selective pressure, TALE expressing ESCs were cultured in the presence of 3 μg/mL blasticidin (Invitrogen Cat# R210-01) and ESC lines stably expressing 3xHA-tagged open reading frames were cultured in the presence of 1 μg/mL puromycin (Sigma-Aldrich, Cat# P8833).

### Generation of the TALE-expressing ESC line

The *Nanog* targeting TALE was created as described in (Briggs et al. 2012) and inserted into the mammalian expression vector pEF-DEST51 (Life Technologies). The vector was then linearized using SgrDI (Thermo Fisher Scientific) and approximately 1.6 × 10^6^ C57BL/6 ESCs were transfected with 15 μg of linearized vector using Lipofectamine 3000 (Thermo Fisher Scientific) according to the manufacturer’s recommendations. As negative control, the cells were transfected with 15 μg of linearized empty pEF-DEST51. 48 h post transfection, blasticidin (Invitrogen Cat# R210-01) was added to the cells a final concentration of 3 μg/mL to eliminate any untransfected cells. To create clonal lines, after approximately two weeks colonies were picked manually and cultured under selective pressure. Lines were then analyzed for TALE expression by western blot using an anti HA-tag antibody (Genscript Cat# A00168-40).

### Generation of ESC lines stably expressing HA-tagged open reading frames

Mouse open reading frame (ORF) expression clones in lentiviral vectors with an EF1α promoter (3xHA-tagged ORF:IRES:mCherry:T2A:puromycin) were obtained for *Sox2* (C-3xHA-tag; United BioResearch Products), *Mybl2* (C-3xHA-tag; VectorBuilder), *Rcor2* (C-3xHA-tag; VectorBuilder), *Wdr76* (N-3xHA tag; VectorBuilder), *Dnmt3l* (C-3xHA-tag; VectorBuilder), *Nfatc2ip* (N-3xHA tag; VectorBuilder) and *Zfp57* (C-3xHA-tag; VectorBuilder). As controls, we utilized two lentiviral vectors with EF1α promoter: “mcherry control” (mCherry:T2A:puromycin) and “3xHA-tag control” (3xHA-tag:IRES:mCherry:T2A:puromycin) (VectorBuilder). Lentiviruses were created by co-transfecting HEK293T cells with the transgene and packaging vectors (PAX2 and MD2.G) as previously described (Larcombe et al. 2019). Briefly, the media was collected at 48 and 72 h post transfection, lentiviral particles were concentrated with PEG precipitation and the titers determined by FACS. C57BL/6 ESCs were transduced with the lentivirus at a multiplicity of infection (MOI) ranging from 0.1 to 1 with 5 μg/μL polybrene (Merck Millipore, Cat# TR1003G). From day three onward (e.g. 48 h post transduction), cells were cultured in the presence of 1 μg/mL puromycin (Sigma-Aldrich, Cat# P8833) to select for transgene integration. Nuclear proteins were isolated using a cell fractionation kit (Cell Signalling Technology Cat# 9038D) and transgene expression was confirmed via western blotting using an anti-HA-tag antibody (Abcam Cat# ab94918).

### Immunofluorescence (IF)

Cultured cells were fixed in 4% paraformaldehyde (PFA) (Sigma Aldrich) in PBS for 10 min and then permeabilized in PBS containing 0.3% Triton X-100 (Sigma Aldrich). Cultures were then incubated with primary antibodies followed by secondary antibodies (see dilutions below). 4’,6-diamidino-2-phenylindole (DAPI) (1:1000) (Life Technologies) was added to visualize cell nuclei. Images were taken with an inverted fluorescence microscope and attached camera (Nikon Eclipse Ti). Primary antibodies used in the study were: anti-HA-tag (Cat# ab94918, Abcam, 1:200), mouse anti-Nanog (Cat# sc-134218, Santa Cruz, 1:100). Secondary antibodies used in the study (all 1:400) were AF488 donkey anti-goat (Cat# A32814, Life Technologies), AF555 donkey anti-mouse IgG (Cat# A31570, Life Technologies).

### Teratoma Assays

Approximately 1.0 × 10^6^ undifferentiated ESCs were injected subcutaneously into the dorsal flanks of immuno-deficient 4 to 20-week-old NOD-SCID mice. Teratomas that developed within 4 weeks post-injection were harvested and fixed in 4% paraformaldehyde (Merck), embedded in paraffin, sectioned at 5 mm and stained with haematoxylin and eosin (H & E). All animal experimentation was performed under the auspices and approval of the Monash University Animal Research Platform animal ethics committee (Approval Number 14351).

### Quantitative RT-PCR

RNA was extracted using the RNeasy Mini Kit (Qiagen) according to the manufacturer’s instructions (with DNase digestion step). RNA was eluted from the columns with 50 μL RNase-free water and quantified using a Nanodrop (Nanodrop Technologies). 500 ng of RNA were then used for cDNA synthesis using the SuperScript III First-Strand Synthesis kit (Invitrogen) according to the manufacturer’s instructions. Real-time quantitative PCR reactions were set up in triplicate with 2 μL of cDNA (1:4 dilution) per reaction using the FastStart Universal SYBR Green Master mix (Rox) (Roche) and run on a 7500 Real-Time PCR System (Applied Biosystems). Gene expression levels were calculated using the 2^−ΔΔCT^ method with *Gapdh* as internal control. The following primer sequences (5’-3’) were used: *Gapdh* (F: AGGTCGGTGTGAACGGATTTG, R: TGTAGACCATGTAGTTGAGGTCA) and *Nanog* (F: TTGCTTACAAGGGTCTGCTACT, R: ACTGGTAGAAGAATCAGGGCT).

### Chromatin Immunoprecipitation qPCR

ChIPs were conducted as described by us previously (Knaupp et al. 2017) with slight modifications. Approximately 30 × 10^6^ cells were used per ChIP. The samples were pre-cleared with 100 μL of Pierce Control Agarose Resin (Thermo Fisher Scientific) for 1 h at 4 °C and immunoprecipitations were performed with 30 μL of Pierce Anti-HA Agarose (Thermo Fisher Scientific) per sample rotating overnight at 4 °C. After DNA purification using the QIAquick PCR Purification Kit (Qiagen Cat# 28106), real-time quantitative PCR reactions were set up in triplicate. 2 μL of DNA were used per reaction with the FastStart Universal SYBR Green Master mix (Rox) (Roche) and run on a 7500 Real-Time PCR System (Applied Biosystems). The percent of input was calculated for each ChIP (% Input = 2^(−ΔCt [normalized ChIP])^). The following primer sequences (5’-3’) were used: *Nanog* promoter (F: AATGAGGTAAAGCCTCTTTTTGG, R: ACCATGGACATTGTAATGCAAA) and background region (e.g. *Sox2* regulatory region 2) (F: GCTAGGCAGGTTCCCCTCTA, R: GCAAGAACTGTCGACTGTGC).

### TALE-mediated Isolation of Nuclear Chromatin (TINC)

Approximately 1 × 10^9^ cells were fixed with formaldehyde as described by us previously (Knaupp et al. 2017) and nuclei and chromatin were isolated according to Kustatscher et al., 2014 to remove non-chromatin-associated proteins. The chromatin was then resuspended in ChIP lysis buffer (1% SDS, 10 mM EDTA, 50 mM Tris pH 8.0) containing protease inhibitor cocktail (Sigma-Aldrich Cat# P3840) at 1g of chromatin per 3.2mL of buffer and sonicated on a Bioruptor NextGen sonication device (Diagenode) to shear the chromatin to an average fragment size of 200 - 500 bp as determined by agarose gel electrophoresis. The samples were then diluted 1:5 with ChIP dilution buffer (165 mM NaCl, 0.01% SDS, 1.1% Triton X-100, 1.2 mM EDTA, 16.7 mM Tris pH 8.0) containing protease inhibitor cocktail (Sigma-Aldrich Cat# P3840) and 0.05% input was collected and put aside on ice. Additionally, 0.05% of the sample was used to determine the DNA concentration to ensure the same amount of chromatin was used for the TALE and the control samples in the TINC assay. To do so, the cross-linking was reversed for 15 min at 100 °C, the sample purified using a QIAquick PCR Purification Kit (Qiagen) and DNA concentration determined using the Qubit dsDNA HS Assay Kit (Thermo Fisher Scientific) and a Qubit 2.0 Fluorometer (Thermo Fisher Scientific). The samples were then precleared with 1 mL of settled Pierce Control Agarose Resin (Thermo Fisher Scientific Cat#26150) for 1h at 4 °C.

TALE-immunoprecipitations were performed with 1 mL of settled Pierce Anti-HA Agarose Resin (Thermo Fisher Scientific Cat# 26182) overnight at 4 °C. Samples were then washed twice for 10 min at 4 °C with dilution buffer containing protease inhibitor cocktail, followed by two washes for 10 min at 4 °C with low salt buffer (150 mM NaCl, 0.5% Na deoxycholate, 0.1% SDS, 1% Nonidet P-40, 1 mM EDTA, 50 mM Tris pH8) containing protease inhibitor cocktail, two washes for 10 min at 4 °C with high salt buffer (500 mM NaCl, 0.5% Na deoxycholate, 0.1% SDS, 1% Nonidet P-40, 1 mM EDTA, 50mM Tris pH8) and a final 5 min wash at 4 °C with TE buffer (0.25 mM EDTA, 10mM Tris pH 8). The volume was 100 mL for each wash step. For elution, the samples were resuspended in 2 mL of 3M NaSCN and incubated rotating for 5 min at room temperature. The eluted samples were removed and the elution step was repeated three times. All four elutions were pooled and concentrated using 3 KDa cut-off concentrators (Amicon Ultra Cat# UFC800308 and UFC500308) to 50 μL. During concentration, the samples were diluted 1:20 with PBS to dilute the NaSCN concentration. 1 μL per sample was then removed and the DNA of this sample as well as of the input samples was then purified as described above to ensure enrichment of the *Nanog* promoter as tested by qPCR as described in the ChIP-qPCR section. The remaining sample was subjected to LC-MS/MS analysis.

### LC-MS/MS analysis

To determine the proteins isolated by TINC, 4 × loading dye (40 % glycerol, 4 % SDS, 400 mM DTT, 0.04 % bromophenol blue, 125 mM Tris pH 8.3) was added to the samples and the samples were incubated at 100°C for 30 min. The samples were then loaded onto a NuPAGE 4-12% Bis-Tris Protein Gel (Thermo Fisher Scientific Cat# NP0335BOX) and a short gel was run using NuPAGE MOPS SDS Running Buffer (Thermo Fisher Scientific Cat# NP0001) to stack the proteins. The proteins were visualized with InstantBlue Coomassie Protein Stain (Expedeon Cat#ISB1L) and the according band excised from the gel. The proteins were reduced with 5 mM DTT (Agilent Technologies Cat# 5188-6439), alkylated with 18 mM 2-chloroacetamide (Sigma-Aldrich Cat# C0267-100G) and in-gel digested overnight at 37°C with sequencing grade trypsin (Promega Cat# V5111). The peptides were then extracted with extraction buffer (50% acetonitrile (Fisher Chemical Cat#A955-4), 5% formic acid (Sigma-Aldrich Cat# 56302-50ML-F)) followed by extraction with 100% acetonitrile. All extracted peptides were combined and dried to completion in an CentriVap Concentrator (Labconco). Samples were then resuspended in 2% acetonitrile, 0.1% formic acid prior to LC-MS/MS analysis.

Using a Dionex UltiMate 3000 RSLCnano system equipped with a Dionex UltiMate 3000 RS autosampler, the samples were loaded via an Acclaim PepMap 100 trap column (100 μm × 2 cm, nanoViper, C18, 5 μm, 100å; Thermo Scientific) onto an Acclaim PepMap RSLC analytical column (75 μm × 50 cm, nanoViper, C18, 2 μm, 100å; Thermo Scientific). The tryptic peptides were separated by increasing concentrations of 80% acetonitrile (CAN) / 0.1% FA at a flow of 250 nl/min for 158 min and analyzed with an Orbitrap Fusion Tribrid mass spectrometer (Thermo Scientific). Each cycle was set to a fixed cycle time of 2 sec consisting of an Orbitrap full ms1 scan (resolution: 120.000; AGC target: 1e6; maximum IT: 54 ms; scan range: 375-1575 m/z) followed by several Orbitrap ms2 scans (resolution: 30.000; AGC target: 2e5; maximum IT: 54 ms; isolation window: 1.4 m/z; HCD Collision Energy: 32%). To minimize repeated sequencing of the same peptides, the dynamic exclusion was set to 15 sec and the ‘exclude isotopes’ option was activated.

Acquired .raw files were searched against the murine UniProtKB/SwissProt database (v2017_07) appended with the TALE protein sequence using Byonic (Protein Metrics) considering a false discovery rate (FDR) of 1% using the target-decoy approach. Carbamidomethylation of cysteine residues was specified as a fixed modification. Oxidation of methionine and acetylation of protein N-termini were selected as variable modifications. Trypsin was used as the enzymatic protease and up to two missed cleavages were allowed. Data visualization and mining was performed in Excel. To minimize false positive protein identifications, we only considered proteins that were identified by at least two unique peptides.

### Data mining

For validation of *Nanog* promoter interactors identified by TINC, ChIP-ATLAS was used for data mining of published ChIP-seq datasets (Oki et al. 2018). Datasets were chosen on the basis of binding between the region 1kb upstream of the *Nanog* enhancer, and the 3’ UTR region of *Nanog*. Furthermore the datasets had to be of untreated/control mouse ESCs. BigWig files and peak files were obtained pre-processed by the ChIP-ATLAS pipeline. The narrow peak files obtained, containing peaks with a false discovery rate cut-off of 0.05. These files were converted into BigBed files using Homer (Heinz et al. 2010). All BigBed and BigWig files were visualised with UCSC mm9 (W. James Kent et al. 2002; W. J. Kent et al. 2010; Raney et al. 2014). For the validation, the data with the following accession numbers was used: SRX2245718, SRX1158299, SRX997153, ERX2244637, SRX039347, SRX4167129, SRX1738884, SRX236477, SRX2441332, SRX1590979, SRX017058, SRX1342343, SRX003864, SRX319555, SRX4394631, SRX476210, SRX2831314, SRX2831316, and SRX017060.

### Enrichment and other statistical analyses

Protein classification analyses were performed with Panther (Thomas et al. 2003), and all other ontology analyses were performed with Metascape (Zhou et al. 2019). Transcription factors were defined by collected annotation from Riken and fantom five mouse TF datasets (Carninci et al. 2005; Kanamori et al. 2004). All other epigenetic modifiers and respective metadata were obtained from the epifactors database (Medvedeva et al. 2015). The pluripotency network was obtained from ESCAPE (Embryonic Stem Cells Atlas of Pluripotency Evidence) (Xu et al. 2013, 2014). Enrichment of TINC proteins within the pluripotency network was calculated with Fisher’s exact test. RNA-seq data of reprogramming fibroblasts into iPSCs was obtained from Knaupp et al. 2017. Raw sequencing files were processed to raw counts as described in the original publication. The raw counts were then TMM normalized and converted to log2 CPM for all analyses. The standardized log2CPM values were centred to the mean expression for each gene. Pearson’s correlation coefficients were calculated on standardized log2CPM values for all genes corresponding to proteins identified in all three TINC replicates. Protein-protein enrichment networks were visualized with Cytoscape v3.7.1 (Shannon et al. 2003). ComplexHeatmap was used for any visualization of heatmaps with default clustering parameters and all other analyses outputs were graphed using GGplot2 unless otherwise stated (Gu, Eils, and Schlesner 2016; Wickham 2016). A set seed of 123 was used in all analyses, unless otherwise stated. Protein-protein interaction network functional enrichment analysis (high confidence (0.700) interaction score) and k-means clustering was conducted using STRING version 11.0 (Szklarczyk et al. 2019). Clusters identified were subjected to gene ontology enrichment using Metascape (Zhou et al. 2019) to identify the most significantly enriched terms for each cluster. Venn diagrams were produced using BioVenn (Hulsen, de Vlieg, and Alkema 2008) or jvenn (Bardou et al. 2014).

## Acknowledgements

This work was supported by the National Health and Medical Research Council (NHMRC) of Australia (GNT1069830) and the Australian Research Council (ARC) Centre of Excellence program in Plant Energy Biology (CE140100008). J.M.P. and R.L. are part of the ARC funded Stem Cells Australia Special Initiative and were supported by Sylvia and Charles Viertel Senior Medical Research Fellowships (J.M.P. and R.L.), NHMRC CDF (J.M.P., APP1036587), ARC Future Fellowship (R.L., FT120100862), and Howard Hughes Medical Institute International Research Scholarship (R.L.). A.S.K. was supported by an NHMRC Early Career Fellowship (APP1092280). We would like to acknowledge the Monash University Histology Platform Paraffin Laboratory for the provision of reagents and technical support.

---

**Supplementary Figure 1.**
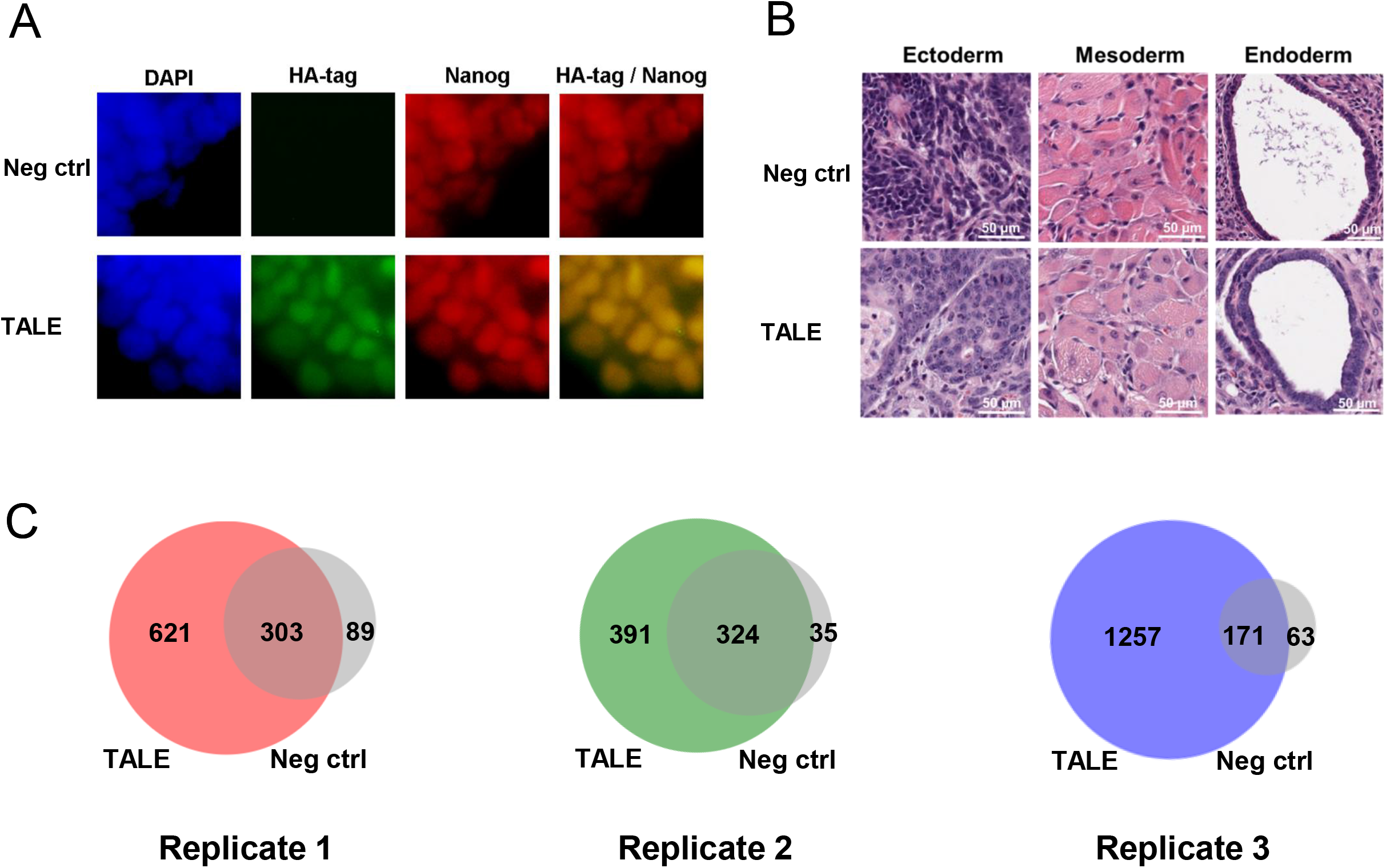
(A) Immunofluorescence shows homogenous expression of the TALE (e.g. anti HA-tag) and unaltered *Nanog* expression. (B) Representative histology sections from teratomas derived from TALE-expressing ESCs and empty vector transfected control ESCs confirming the presence of derivatives of all three germ layers. (C) Overlap of the proteins identified in each TALE replicate and the according negative control. Only proteins solely identified in the TALE sample but not the negative control were utilized for further analysis.

**Supplementary Figure 2.**
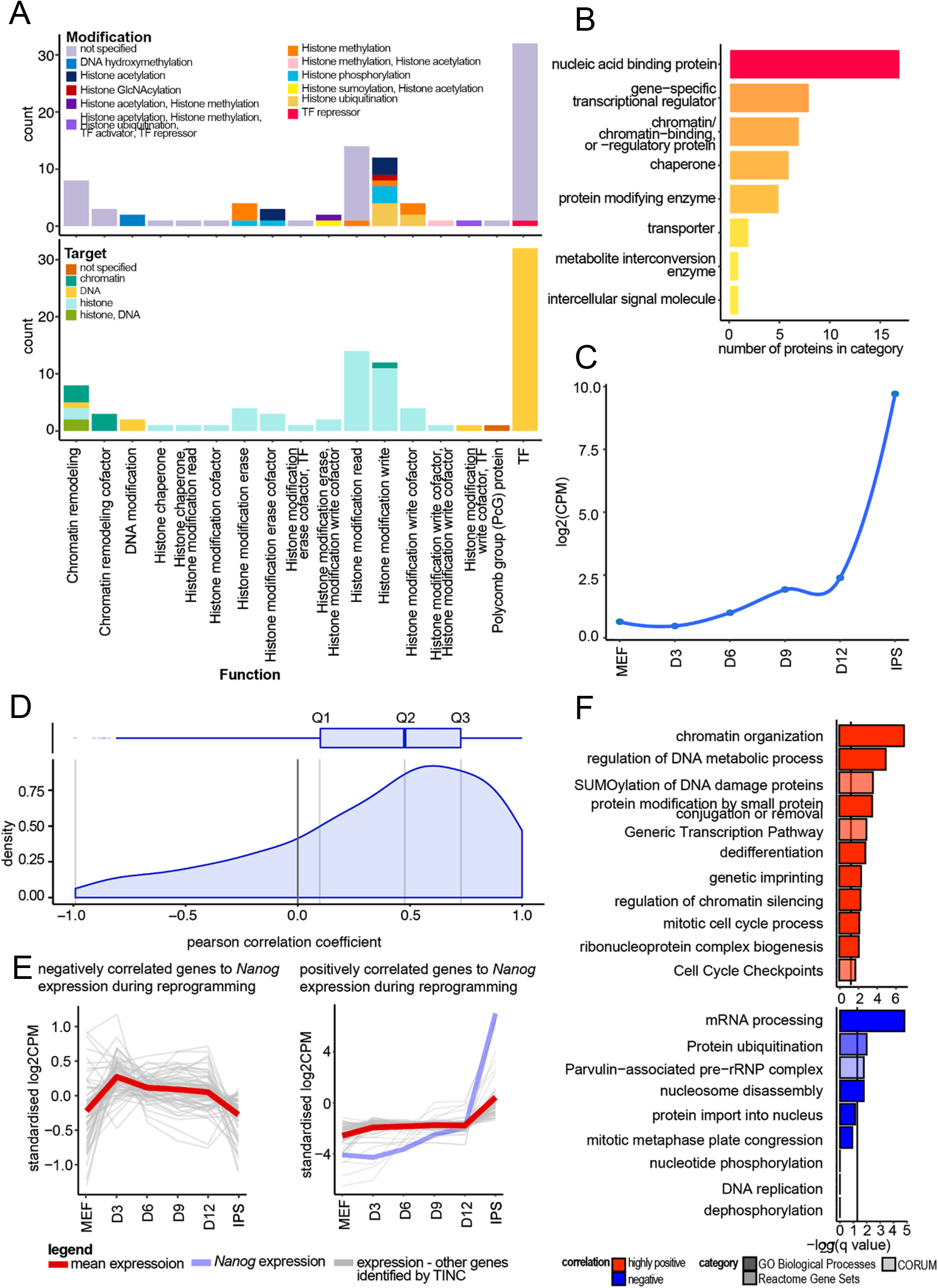
(A) The classification of all TFs and epigenetic modifiers identified by TINC at the *Nanog* promoter. The epigenetic factors were first categorized by their role or function and then further classified by the type of modification they exert as well as the type of molecules they target. (B) Panther protein classification of the TINC proteins identified as part of the pluripotency regulatory network. The color gradient corresponds to the number of proteins in the term with yellow being a few and red being many. (C) The log2CPM expression of *Nanog* throughout reprogramming. (D) Distribution of Pearson’s correlation coefficients of the expression of the candidates identified by TINC in relation to the expression of *Nanog* during reprogramming of MEFs to iPSCs. (E) The standardized expression values of all negatively (62) and positively (218) correlated genes in comparison to *Nanog* during reprogramming of MEFs to iPSCs. Shown are also the mean trends of these TINC proteins during reprogramming as well as the expression of *Nanog*. (F) Gene ontology enrichment of biological processes and reactome gene sets of proteins identified by TINC at the *Nanog* promoter highly positively (top quantile, 70 genes) and negatively correlated to *Nanog* expression during reprogramming.

